# Verification of CRISPR editing and finding transgenic inserts by Xdrop™ Indirect sequence capture followed by short- and long- read sequencing

**DOI:** 10.1101/2020.05.28.105718

**Authors:** Blondal Thorarinn, Gamba Cristina, Jagd Lea Møller, Su Ling, Demirov Dimiter, Guo Shuang, Camille M. Johnston, Eva M. Riising, Wu Xiaolin, Marie J. Mikkelsen, Szabova Ludmila, Mouritzen Peter

## Abstract

Validation of CRISPR-Cas9 editing typically explore the immediate vicinity of the gene editing site and distal off-target sequences, which have led to the conclusion that CRISPR-Cas9 editing is very specific. However, an increasing number of studies suggest that on-target unintended editing events like deletions and insertions are relatively frequent but unfortunately often missed in the validation of CRISPR-Cas9 editing. The deletions may be several kilobases-long and only affect one allele. The gold standard in molecular validation of gene editing is direct sequencing of relatively short PCR amplicons. This approach allows the detection of small editing events but fails in detecting large rearrangements, in particular when only one allele is affected. Detection of large rearrangements requires that an extended region is analyzed and the characterization of events may benefit from long-read sequencing. Here we implemented Xdrop™, a new microfluidic technology that allows targeted enrichment of long regions (~ 100 kb) using just a single standard PCR primer set. Sequencing of the enriched CRISPR-Cas9 gene edited region in 4 cell lines on long- and short -read sequencing platforms unravelled unknown and unintended genome editing events. The analysis revealed accidental kb large insertions in 3 of the cell lines, which remained undetected using standard procedures. We also applied the targeted enrichment approach to identify the integration site of a transgene in a mouse line. The results demonstrate the potential of this technology in gene editing validation as well as in more classic transgenics.

## 1. Introduction

The high precision, specificity and efficiency of CRISPR has provided an unprecedented improvement in genetic engineering with a performance spectacularly improved relative to former technologies for gene targeting [1,2]. The advancement has accelerated an extended exploitation of genetic engineering in organisms from plants and animals to humans. This and considerations of applying CRISPR for gene therapy in humans have spurred concerns about safety where the most pressing questions currently relate to accuracy of the applied CRISPR editing and risk of potential off-target editing.

The concern about distal off-target events understandably relates to the potential consequences of undesired interruption of function for non-target genes elsewhere in the genome - in particular in genomic regions containing oncogenes and tumor suppressor genes [3]. A recent review of studies applying CRISPR-Cas9 in mouse and human cells suggests that distal off-target events are relatively rare and undetectable [4]. While this type of distal off-target events may be undetectable, other unintended editing events are occasionally detected including duplications, indels, single nucleotide variants (SNV), and also larger rearrangements [5]. Reintegration of larger excised regions in the vicinity of the edited region may occur when applying strategies using two gRNAs simultaneously [6].

However, strategies applying just a single gRNA may also produce unintended editing in the form of larger deletions and smaller indels [7]. Focused analysis directed towards a more extended region surrounding the site of gene editing suggests that on-target unintended editing may in fact not be infrequent [8]. The unintended editing events occur during repair of the Cas-9 nuclease generated double strand breaks (DSB) by either the non-homologous end-joining (NHEJ) repair pathway or homology-directed repair (HDR) pathway [9].

A multitude of methods are employed to verify correct gene editing and to ensure that on-target unintended editing events have not occurred. The most commonly applied molecular method is direct DNA sequencing by Sanger or NGS of PCR amplicons, which are typically < 1000 bp in size [10–12]. Other common methodologies include T7 endonuclease 1 (T7E1) mismatch detection assay, Tracking of Indels by Decomposition (TIDE) assay and the Indel Detection by Amplicon Analysis (IDAA), cloning of PCR amplicons followed by sequencing, but occasionally also WGS, array-Comparative Genomic Hybridization (CGH), Southern blot, Fiber-FISH and FISH [6,7,9]. Mostly these methods just provide information from a limited region around the gene editing site and will not capture the entire sequences from CRISPR/Cas mutagenesis effects and in particular not detect larger deletions [13,14]. The work by Kosicki et al 2018 [8] have underlined the importance of extending the molecular analysis to include several kilobases surrounding the site of CRISPR-Cas9 induced DSB and demonstrates how long-read sequencing may ease the resolution of occurred rearrangements.

Here we describe the application of a new indirect sequence capture technology (Xdrop™) for enrichment of long genomic DNA fragments to analyse genotypes and allele status in CRISPR-Cas9 edited cells by both long- and short-read sequencing. Surprisingly, we detect CRISPR-Cas9 induced unintended gene editing in a set of CRISPR modified induced pluripotent stem (iPS) cell lines. Earlier attempts to validate the performed CRISPR editing did not detect these unintended edits despite comprehensive analyses using several complementary approaches [15]. We also apply the Indirect sequence capture technology to identify the integration site of a transgene in a mouse line generated with more classical zygote transformation procedures.

### 1.1 Description of the technology

The Xdrop™ indirect sequence capture relies on partitioning of HMW DNA into millions of picolitre reactions inside double emulsion droplet compartments along with PCR reagents and primers to amplify a single small (~150 bp) amplicon (“detection sequence”) located within the region of interest. Indirect sequence capture refers to the location of the detection sequence which may be located at a distance 5-10 Kb or more from the main region of interest such as the site of gene editing. Droplets containing the DNA of interest are identified by droplet PCR (Fig.1) followed by staining with a DNA intercalating fluorescent dye. Only droplets containing DNA from the region of interest (ROI) will generate a detection sequence amplicon and become fluorescent. Because of the size and composition of the double emulsion droplets, these can be sorted on a standard flow cytometry cell sorter to enrich for fluorescent droplets. The sorted droplets containing long DNA fragments are then isolated and amplified by single molecule multiple displacement amplification in droplets (dMDA). The amplification capacity of the dMDA is able to generate ~1.5 µg of amplified DNA from 6 pg of input DNA and will be most efficient on molecules larger than 5 kb. The resulting enriched DNA is compatible with both short- and long-read sequencing platforms.

The size of the enriched region is proportional to the integrity of the input DNA. Input DNA with a size of ∼50 kb will theoretically result in enrichment of a region no larger than a 100 kb. Enrichment of a large region will therefore be inefficient with degraded DNA but may be sufficient for a short region of a few kb in size. The enrichment only requires the design of a single primer pair to generate the short detection sequence amplicon that is exclusively used for detection, selection and enrichment of the ROI. The amplicon is rarely detected in downstream sequencing because of poor amplification of short molecules by the dMDA and because of small DNA molecule losses during droplet breaking and size selections applied during sequencing library preparation procedures.

**Figure 1.**
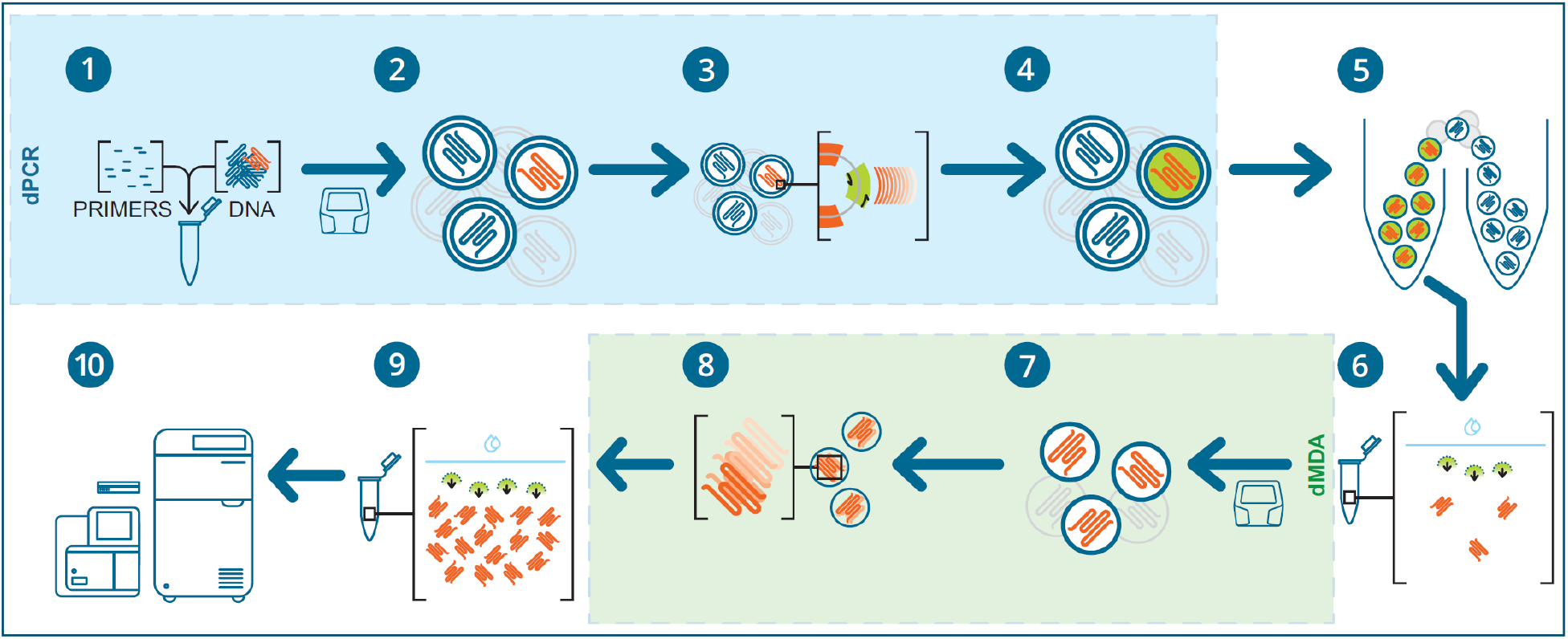
Indirect sequence capture and unbiased amplification with Xdrop™. The workflow includes indirect sequence capture in double emulsion droplets (dPCR box, blue) and multiple displacement amplification in single emulsion droplets (dMDA box, green). In the first step (1), the sample is mixed with the Detection Primers and PCR reagents and this mix (2) is partitioned into millions of double emulsion droplets of ~20 µm diameter using the Xdrop™ instrument and the dPCR cartridge. These droplets are highly stable and are suitable for standard PCR cycling, flow cytometry analysis, and sorting. (3) Droplets containing the Region of Interest (ROI) are identified by the detection sequence, a short amplicon (∼150 bp) placed within or adjacent to the ROI. Droplets are stained with an intercalating dye (4) and positive droplets are (5) identified by their fluorescence and physically separated from negative droplets using a standard cell sorter. DNA is released from double emulsion droplets (6) resulting in a population of long DNA molecules enriched for the ROI and comprising kilobases of information. For downstream DNA amplification, (7) each long fragment derived from the enrichment is partitioned as single molecules into thousands of single emulsion ∼85 µm diameter droplets for (8) high fidelity multiple displacement amplification in droplets (dMDA). Amplified enriched fragments from the ROI are then (9) released from the single emulsion droplets. The enriched DNA is compatible with (10) downstream analyses, such as long- and short-read sequencing.

## 2. Material and methods

### 2.1 Samples

#### iPS cell lines

The iPSC line BIONi010-C and four CRISPR-Cas9 gene edited *APOE* derivatives were described in [15]. Three of these cell lines were edited at single nucleotide positions in Exon 4 to modify genotypes at two different SNPs (rs429358 and rs7412), recognized as the most common genetic risk factor for sporadic Alzheimer’s disease (AD). The editing was performed to obtain isogenic cell lines homozygous at the two SNP positions namely *APOE*-ε4/ε4 (rs429358 C/C, rs7412 C/C), *APOE*-ε3/ε3 (rs429358 T/T, rs7412 C/C) and *APOE*-ε2/ε2 (rs429358 T/T, rs7412 T/T). The *APOE*-ε2 (rs429358 T, rs7412 T) and *APOE*-ε4 (rs429358 C, rs7412 C) haplotypes are associated with lower and higher risk of AD, respectively [16,17]. The cell lines were developed to serve as an isogenic AD model system to be able to study the biological mechanisms by which the *APOE* alleles affect the risk of developing AD [15]. Moreover, an *APOE* knock-out cell line was generated, carrying an 8-nucleotide deletion in Exon 2. All cell lines were generated by editing the iPSC line BIONi010-C (genotype *APOE*-ε3/ε4), previously generated from human skin fibroblasts [18], apart from *APOE*-ε2/ε2 which was generated from the *APOE*-ε3/ε3 cell line [15].

We analysed the *APOE* locus and surrounding region in all five cell lines.

#### Pax8-CreERT2 Transgenic mouse line

To generate Pax8-CreERT2 transgene (Supplementary Fig 1), a plasmid containing Pax8-rtTA sequence was obtained from Dr. Robert Koesters [19] from INSERM/Université Pierre et Marie Curie, France. To make this transgene tamoxifen-inducible rtTA sequence was swapped for CreERT2 sequence from pCAG-CreERT2 plasmid (Addgene). Here the gene for Cre recombinase is fused with a mutated ligand-binding domain of the estrogen receptor (ER), thus allowing Cre activity be controlled by tamoxifen. The ER domain sequesters the Cre in the cytoplasm until tamoxifen binds to the ER and the complex translocates to the cell nucleus where Cre recombines loxP sites of the modified genes (floxed genes). The Pax8-CreERT2 transgene was linearized using SalI and NotI restriction enzyme digest and the resulting ∼8.4 kb Pax8-CreERT2 DNA fragment was microinjected, using standard procedure, into the pronucleus of fertilized mouse eggs of C57Bl/6 mice (The Jackson Laboratory) to generate transgenic founders. Founders were bred to C57Bl/6 mice (The Jackson Laboratory) to observe transgene transmission to next generations. Transgenic lines that transmitted the transgene were checked for the transgene expression by mating to ROSA26R reporter mouse (The Jackson Laboratory) and by evaluating the offspring for expression of beta galactosidase by histological staining. One line with desired expression pattern was selected for analysis.

### 2.2 Target enrichment by Xdrop™

#### 2.2.1 Input DNA

High molecular weight (HMW) DNA was isolated from thawed *APOE* cell lines by Bioneer A/S using the Blood & Cell Culture DNA Mini Kit (Qiagen). For analysis of *Pax8-CreERT2* insert high molecular DNA was prepared from mouse ears using MagAttract HMW DNA Kit (Qiagen) following the manufacturer’s protocol. Around 300 ng of the isolated genomic DNA were further subjected to capture bead clean-up step utilizing HighPrep™ PCR Clean-up Capture bead system (MagBio Genomics Inc.) in 1:1 (Vol:Vol) ratio according to the manufacturer’s instructions. HMW DNA was resuspended in 10 mM Tris pH 8 and quantified by Quantus™ Fluorometer (Promega Inc). The DNA size and integrity were assessed on TapeStation™ instrument (Agilent, Genomic DNA ScreenTape), and all samples had DNA > 50 kb in size. For *APOE* cell lines 10 ng and for *Pax8-CreERT2* mouse 5 ng of HMW DNA was used as input to the Xdrop™ workflow.

In order to obtain a significant enrichment, we calculated the optimal amount of input template DNA using the automated online tool “DNA input calculator” available online at Xdrop homepage.

#### 2.2.2 Detection sequence design and testing

Using the “Primer design tool” available online at the Xdrop homepage we designed dPCR and qPCR primer sets. The primer tool will check for primer specificity in the genome but this is also tested by qPCR (see below). For *APOE* we designed the detection sequence primer set within *APOE* intron 1 ~ 2kb upstream the CRISPR-Cas9 edited region (fw: GCTAGCCGTCGATTGGAGAA, rev: CATCTCAGTCCCAGTCTCGC, amplicon position: chr19: 44,906,188 – 44,906,317) see Supplementary Fig. 1. We also designed a second primer set, the validation primer set ∼2.5kb away from the detection sequence (fw: GGAGGCCTCCGTTTTCTCAA, rev: CCGGACTTAAGGCAGCATCA, amplicon position: chr19: 44,903,692 – 44,903,842) to validate the enrichment of the ROI by qPCR at the end of the Xdrop™ workflow. For the *Pax8-CreERT2* insert enrichment, the primer sets for detection and validation sequences were designed within the human estrogen receptor sequence to be specific for the insert itself see Supplementary Fig. 2 (detection sequence primer set; Fwd: ATGATTGGTCTCGTCTGGCG, Rev: ATGCGGAACCGAGATGATGT, (validation primer set; fwd: ACAGGGAGCTGGTTCACATG, rev: TCAGGATCTCTAGCCAGGCA).

Both Detection and Validation Primers were tested prior to the enrichment by qPCR, using as input a representative HMW DNA sample and the Primer test PCR kit (Samplix ApS), following manufacturer’s recommendations. Primer set approval criteria were sigmoid shaped amplification curves with PCR amplification efficiencies between 85-115% to ensure good signal generation in droplets. Efficiency was calculated based on 10-fold dilution series of input DNA (0.4 ng, 4 ng, 40 ng) in triplicate measurements. To evaluate specificity amplicon melting profiles were generated and evaluated for single peak amplicon melting profiles to ensure absence of primer dimer formation or any off-target amplification. The specificity is of utmost importance to avoid sorting of false positive droplets with non-target DNA as this will compromise enrichment.

#### 2.2.3 Target detection by dPCR

Double emulsion droplets were generated in the Xdrop™ instrument by using the dPCR cartridge and the dPCR kit (Samplix ApS). The dPCR cartridge was loaded with the 1x dPCR buffer, dPCR mix including detection sequence primers and 5 or 10 ng HMW DNA, and dPCR oil in the mentioned order to ensure successful droplet production and sealed with the rubber gasket following manufacturer’s recommendations.

After droplet generation, droplets were collected, aliquoted in four 0.2 mL Forensic DNA Grade PCR Tubes, (Eppendorf) and transferred to a S1000 thermal cycler (Bio-Rad Laboratories), where the following program with slow ramping at all stages (0.5°C/sec) was run: 30°C (5 sec), 94°C (3 min), 40 cycles of 94°C (3 sec) and 60°C (30 sec), 4°C (∞). Droplets were stored overnight at 4°C before flow cytometer sorting.

#### 2.2.4 Flow cytometry sorting of target

A Sony SH800S cell sorter with 100 μm nozzle sorting chip (Sony Biotechnology) was used to sort the positive dPCR droplets (containing long DNA fragments from the ROI) from the negative droplet population. The cell sorter was equipped with a blue laser (488 nm) and optical configuration detecting events in the green spectrum (optical filter pattern 1, FL1). The dPCR droplets were stained with the intercalating fluorescent Droplet dye (Samplix ApS) before sorting following manufacturer’s instructions. The Cell sorter control kit (Samplix ApS), consisting of ready-made dPCR droplets with a large population of positive droplets (10-15% of total dPCR droplets), was used to ease pre-setting the flow cytometry sorting gates using a sample pressure set to aim for ~ 5000 events/sec. Positive droplets were gated and sorted out into 15 µL of PCR grade water in a 1.5 ml DNA LoBind tube (Eppendorf).

The observed number of positive droplets correlated with the theoretical number calculated using the “Enrichment Predictor” tool available online at Xdrop homepage (see Table 1). After sorting, the positively sorted dPCR droplets were kept for a short time at 4°C before dMDA amplification. To prevent degradation and loss of sorted long DNA fragments, the dMDA was always performed on the same day as the droplet sorting.

#### 2.2.5 Droplet multiple displacement amplification (dMDA)

After sorting, the DNA was released from the positive droplets by adding Break Solution and Break colour (Samplix ApS) to each sample. After brief centrifugation, all of the clear bottom phase (Break solution) was carefully collected and discarded. The upper phase containing the enriched DNA was used to set up dMDA reactions using the dMDA kit (Samplix ApS) and following manufacturer’s recommendations. To monitor contamination, we included as negative controls both aliquots of sheath fluid from the flow cytometer and PCR grade water. As a positive control we used 1 pg Human Genomic DNA, Female (Promega Corp.). The samples and the dMDA reagents were mixed and injected into the dMDA cartridge followed by overlaying with dMDA oil. The loading of reagents in the dMDA cartridge carefully followed manufacturer’s recommendations. The gasket sealed cartridge was inserted into the Xdrop™ instrument for single emulsion droplet generation. Then droplets were collected and transferred into a 0.2 mL Forensic DNA Grade PCR Tube (Eppendorf) and incubated in a thermal cycler for 16 hours at 30°C, where the reaction was terminated at 65°C (10 min) and then kept at 4°C until harvested.

After dMDA incubation, DNA was harvested by adding Break Solution and Break colour from the dMDA kit (Samplix ApS) to each sample, followed by pipetting and discarding the clear bottom phase (Break solution).

#### 2.2.6 Evaluation of enrichment

The total amount of enriched DNA released from the dMDA droplets was measured with Quantus™ Fluorometer (Promega Inc.) and fragment size distribution was inspected on TapeStation™ (Agilent, Genomic DNA ScreenTape), following manufacturer’s instructions. Fold enrichment of target DNA was firstly assessed by qPCR using the Primer test PCR kit (Samplix ApS) and the validation primers, not overlapping with the detection sequence. Assessment by qPCR was performed following manufacturer’s instructions to provide a rough estimate of the enrichment success prior to sequencing. The cut-off applied here was at least 100 fold enrichment measured by qPCR and all samples complied with this criteria. Calculation of fold enrichment was assessed with the online “Enrichment calculator” available at Xdrop homepage.

#### 2.2.7 Insert confirmation

##### iPSC lines

The breakpoints and inserts found in the ε2/ε2 and ε3/ε3 cell lines was identified by long- and short-read sequencing of the enriched sample, but also further confirmed by standard PCR coupled with short-read sequencing. Four PCR assays were designed which in combination cover the identified left and right breakpoints between the 3.4 kb insert and the *APOE* genomic locus sequences (forward primer in *APOE* gene and reverse primer in insert for the left breakpoint and vice-versa for the right breakpoint. PCR products were generated and submitted for short-read sequencing as described in 2.3.1. Primer pairs used were Assay 1 fwd: CCGTTCCTTCTCTCCCTCTT, Assay 1 rev: GGCATCCCAGAAGTGTGAGT, Assay 2 fwd: AGGGAACAAAAGCTGTCGAG, Assay 2 rev: CGATATCGAATTCAAGCTTTCTA, Assay 3_fwd: CGCATGTCACTCATCGAAAG, Assay 3 rev: ACAGTGGGAGTGGCACCTT, Assay 4 fwd: GATCGGCCATTGAACAAGAT and Assay 4 rev: GGCGTTCAGTGATTGTCG. The four PCR assays were run on a thermal cycler using 10 ng DNA from the ε2/ε2 and ε3/ε3 cell lines with PrimeStar GXL PCR mastermix (Takara Bio) according to the manufacturer’s instruction using 58 °C annealing temperature. PCR products were evaluated and quantified by Tapestation™ System (Agilent, D5000 ScreenTape) according to the manufacturer’s instructions and 200 ng of each PCR pooled, followed by bead purification using HighPrep™ PCR Clean-up Capture bead system (MagBio Genomics Inc.) in 1:1.8 (Vol:Vol) ratio according to the manufacturer’s instructions. DNA was resuspended in 30 uL PCR grade water and quantified by Quantus™ Fluorometer (Promega Inc). The PCR product pools were sequenced by short read sequencing as described by 2.3.1.

##### Pax8-CreERT2 Transgenic mouse line

In order to confirm the borders of the insertion of the *Pax8-CreERT2* construct into chromosome 1, standard PCR and Sanger sequencing was performed. Two PCR assays for each border was used. Left border Assay 1 fwd: CTGAGGGTTTCCCATTCAGCA, rev: CCGATCTGCTCACTCGCAG, Assay 2 fwd: GGGGCCTAGAAAGATGTATGGT, rev: TCAGGGCTTCCACTAGCAC. Right border: Assay 1 fwd: GTAGTGACCCTGGCCTTGTA, rev: GTTCTGTTGCTGCTGGCATTA, Assay 2 fwd: GCACCTACCATTCTCCCACTA, rev: GCTCTTGTCATTTCTCATCAGGG. The four PCR assays were run on a thermal cycler using 100 ng DNA with PrimeStar GXL PCR mastermix (Takara Bio) according to the manufacturer’s instruction using 58 °C annealing temperature. PCR products were evaluated and quantified by Tapestation™ System (Agilent, D5000 ScreenTape) according to the manufacturer’s instructions. The amplicons were Sanger sequenced at Eurofins Genomics.

### 2.3 Library preparation and sequencing

#### 2.3.1 Short-read sequencing

For short-read sequencing library generation, 300 ng of enriched DNA samples were fragmented using the NEBNext® Ultra™ II FS DNA Library Prep Kit (New England Biolabs) according to the manufacturer’s recommendation with 5 minutes incubation time. After fragmentation, the samples were processed according to the TruSeq PCR Free Library Preparation protocol starting with end-repair, size selection, followed by dA tailing, barcoding and adaptor ligation as described by the manufacturer (Illumina Inc.). The size distribution of libraries was evaluated by Tapestation™ System (Agilent, D1000 ScreenTape) according to the manufacturer’s instructions. Concentrations were calculated and the samples pooled in equimolar concentrations. Finally, the sample pool was quantified using KAPA Library Quantification Kit for Illumina and BioRad qPCR platforms (Roche) according to manufacturer’s instructions. Short-read sequencing was performed on iSEQ100 desktop sequencer according to the manufacturer’s instructions (Illumina Inc.) using paired end 2 x 151 protocol. Basecalling was performed automatically using GenerateFASTQ - 2.0.0 module.

#### 2.3.2 Long-read sequencing

For long-read sequencing library generation, 1.5 µg of enriched DNA samples from the Xdrop™ workflow were used to generate Oxford Nanopore long-read sequencing libraries. First the amplified DNA was subjected to debranching using T7 endonuclease I. This is crucial when working with MDA amplified DNA as the branched DNA rapidly will block the nanopores. Apart from cleaving non-β DNA structures like branched DNA, T7 endonuclease I will cut at DNA mismatches. Debranching was performed by incubation with 15 units of T7 endonuclease I (New England Biolabs) for 15 minutes at 37 °C. The DNA samples were DNA repaired and end-prepped using the NEBNext FFPE DNA Repair Mix (New England Biolabs) and NEBNext End repair/dA-tailing Module (New England Biolabs), and then barcoded using the Native Barcoding kit (Oxford Nanopore Technologies) followed by adaptor ligation by the Ligation Sequencing kit (Oxford Nanopore Technologies). All procedures were carried out following manufacturer’s instructions and Samplix recommendations for Xdrop enriched DNA. Freshly prepared libraries were quantified by Quantus™ Fluorometer (Promega Corp.) according the manufacturer’s recommendations and 30-50 fmol DNA loaded into a MinION flow cell (R9.4.1, Oxford Nanopore Technologies) and run on standard setting for 16 hours. Base calling of native reads was performed using Guppy v. 3.4.5 with high accuracy and quality filtering (Oxford Nanopore).

### 2.4 Data analysis

#### 2.4.1 Long-read data

For data from *APOE* cell lines mapping was performed using Minimap 2.0 [20] with default settings against the human genome GRCh38.1, as well as the reconstructed hybrid sequence of the human genome (including the unintended insertion from the modified pEasyFlox plasmid). Coverage was calculated using SAMtools [21] and BEDTools [22], and visualized in R [23]. Mapped reads were visualised using IGV 2.8.2 [24].

For data from *Pax8-CreERT2* mouse lines mapping was performed using Minimap 2.0 [17] with default settings against the *Pax8-CreERT2* construct. The reads mapping to the construct was extracted using SAMtools [21] and mapped using Minimap2.0 against the mouse reference genome GCF_000001635.26_GRCm38.p6. Coverage and reads were inspected using CLC genomics Workbench (Qiagen), and a new reference genome with the inserts were build All reads were mapped to the GCF_000001635.26_GRCm38.p6 with the inserts using Minimap 2.0 [27]. Coverage was calculated from the different bam files using SAMtools [21] and BEDTools [19].

#### 2.4.2 Short-reads data

Illumina reads were trimmed with Trimmomatic [25] using default settings and mapped to the construct and the human genome GRCh38.1 with BWA-MEM [21]. Coverage was calculated using SAMtools and BEDTools [22], and visualized in R [23].

#### 2.4.3 Enrichment calculation

Sequencing-based enrichment was calculated for both short- and long-reads data using the following equation: Enrichment = (number of target reads / total mapped reads to reference) - (Target size in bp / reference genome size in bp) = (proportion of reads mapped to target region) / (proportion of target region to reference).

## 3. Results

### 3.1 Enrichment of edited *APOE* gene region and of Pax8-CreERT2 transgene

The samples selected for this study was chosen to investigate whether the indirect sequence capture technology could be applied to validate CRISPR editing site and to locate the integration site of a transgene. For the former we chose the Alzheimer’s disease model which was obtained from Bioneer A/S upon consultation with Dr. Bjørn Holst. For the latter we used a genetically engineered mouse line which had already been developed at the Center for Advanced Preclinical Research at NCI.

The droplet generation and droplet PCR were performed after assay validation by qPCR showed that efficiency of the assays was good (PCR efficiency about 100% ± 15 %) with single-peak amplicon melting profiles indicating high PCR specificity, and no-template controls without signal showing absence of primer dimers. The high efficiency is required to produce good signal in positive droplets relative to negative droplets containing non-targets. In the optimal situation the population of positive droplets should form a distinct cluster separated from the negative droplets (see Fig. 2a). The high specificity is important as any false positive droplet will contain non-target DNA which will affect enrichment negatively. Droplet generation and droplet sorting were successful for all samples. Overall, the positive droplets represented around 0.01 % of the total double emulsion droplet population. We sorted enough double emulsion droplets to get 300-600 positive droplets from each droplet production (see Table 1). An example of the gating strategy is shown in Fig. 2a for the droplet sorting for sample *APOE*-ε3/ε3. Here a total 4.29 million double emulsion droplets were detected resulting in 467 positive droplets sorted.

**Table 1.** Summary of flow cytometry sorting statistics for the Pax8-CreERT2 transgenic mouse line and the five iPSC lines including four CRISPR edited lines. Input DNA: the amount of genomic DNA used as input for the Xdrop enrichment. No. of dPCR droplets analysed: total double emulsion droplets analysed by flow cytometry for each sample. No. of positive droplets sorted: number of double emulsion droplets amplifying the Detection Sequence and sorted by flow cytometry. % positive sorted to total analysed: double emulsion droplets identified as containing the target sequence compared to the total amount of double emulsion droplets analysed. % positive droplets predicted: Expected percentage of double emulsion droplets containing the target sequence considering the amount of input DNA, size of target genome and targets per genome.

| Sample | Input DNA | No. dPCR droplets analyzed | No. of positive droplets sorted | % positive sorted to total analyzed | % positive droplets predicted | DNA after dMDA (ng) | Enrichment measured by qPCR |
| --- | --- | --- | --- | --- | --- | --- | --- |
| *Pax8-CreERT2 mouse* | 5 ng | 3.58 x 10<sup>6</sup> | 448 | 0.013% | 0.008% | 675 | >100x |
| *APOE-KO* | 10 ng | 3.66 x 10<sup>6</sup> | 477 | 0.013% | 0.012% | 825 | 350x |
| *APOE*-ε4/ε4 | 10 ng | 4.13 x 10<sup>6</sup> | 554 | 0.013% | 0.012% | 810 | 1890x |
| *APOE*-ε2/ε2 | 10 ng | 4.08 x 10<sup>6</sup> | 498 | 0.012% | 0.012% | 960 | 1889x |
| *APOE*-ε3/ε4 | 10 ng | 4.28 x 10<sup>6</sup> | 335 | 0.008% | 0.012% | 1067 | 3697x |
| *APOE*-ε3/ε3 | 10 ng | 4.29 x 10<sup>6</sup> | 467 | 0.011% | 0.012% | 1050 | 1248x |

**Figure 2.**
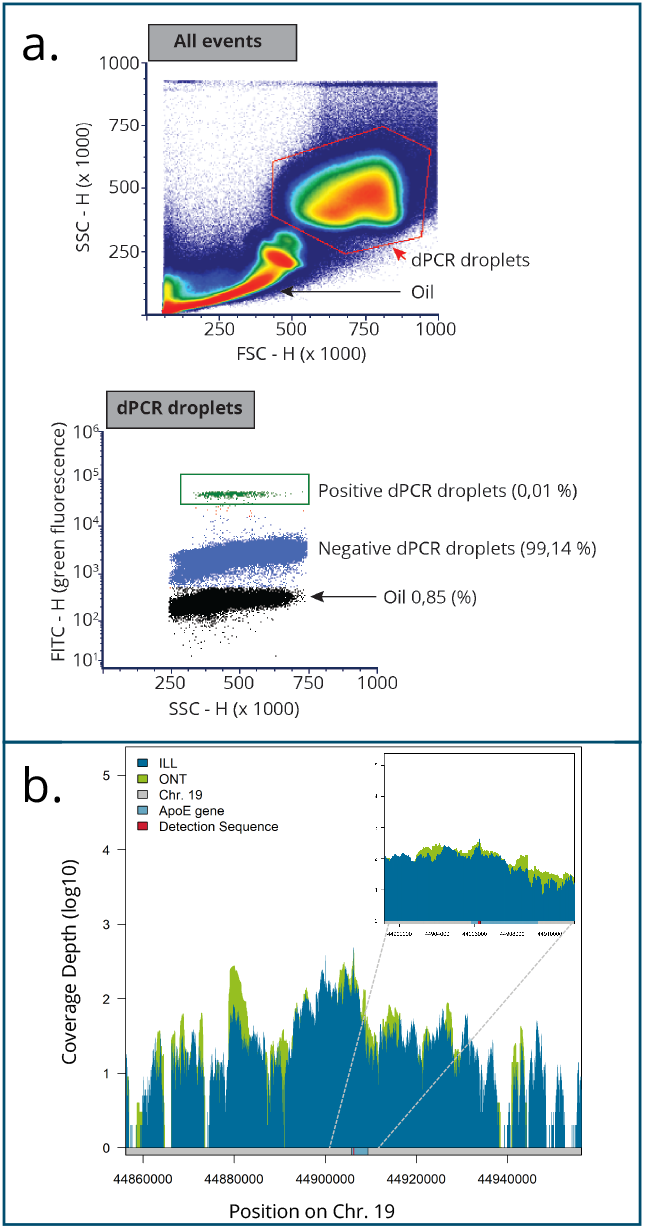
Sorting of dPCR droplets and sequencing coverage of the enriched target. a) top, First, we identify the double emulsion dPCR droplets by plotting forward scatter (FSC) versus side scatter (SSC) using the “height” setting, which allows identifying the population of dPCR droplets here indicated by the polygonal red frame. All events from the sort are shown. bottom, Second, after gating the dPCR droplets, we identify the positive droplets based on the green fluorescent signal (FITC) versus SSC. Based on this gating strategy, the positive population of dPCR droplets is collected (green gate rectangle). Thus, a negative and positive external control is not required since the sample is not stained with an antibody and since an internal control is present. The sorting gates can be moved during sorting to capture as many positive dPCR droplets as possible. b) Depth of coverage of Illumina (blue) and Oxford Nanopore (green) sequencing data (sample ε2/ε2). The main graph focuses on the 100 kb surrounding the detection sequence (lower bar, red) designed to capture the APOE gene region (lower bar, light blue). On the top right corner, a zoomed in area on the central 10 kb surrounding the detection sequence.

Long DNA fragments were isolated from the positive sorted droplets and amplified in single emulsion droplets (dMDAs). The Xdrop™ workflow provided approximately 1.5 µg of enriched DNA per sample, starting from 5 or 10 ng of HMW DNA. Evaluation of enrichment by qPCR estimated an enrichment of 1000-1800 fold which was well above the cut-off criteria of 100 fold. After enrichment, we proceeded with both long- and short-read sequencing (see Fig. 2b), on Oxford Nanopore (samples ε3/ε3, ε2/ε2 and the mouse line) and Illumina (all iPSC samples). Approximately 150,000 long reads (Oxford Nanopore) were obtained per sample (Supplementary Table 1) and enrichment estimations on this data was ∼1000-fold for the *APOE* gene and ∼8000-fold for the *Pax8-CreERT2* transgene insert (see Table 2). Approximately 5 million paired-end reads of 151 bp (Illumina) were obtained per sample and enrichment estimations on this data was ~700-fold for the *APOE* gene (see Table 2), in overall agreement with Oxford Nanopore data.

**Table 2.**
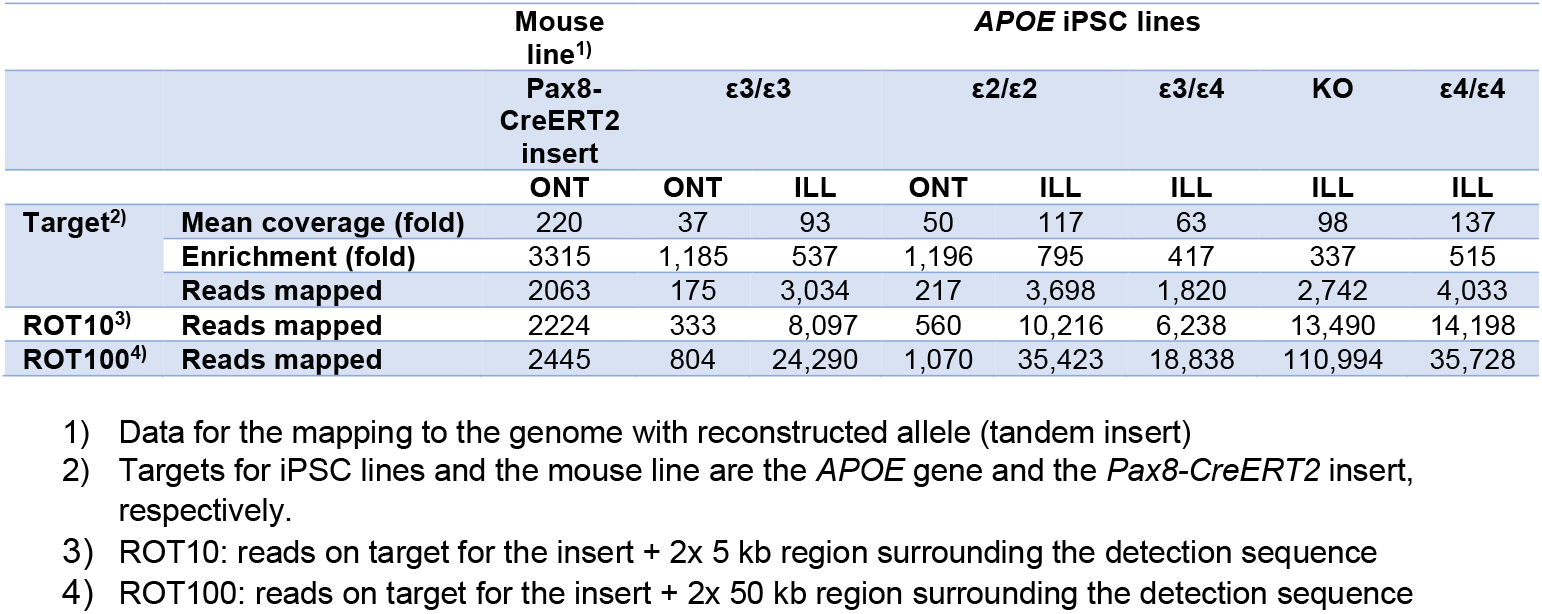
Sequencing read summary for mouse line and *APOE* iPSC lines.

### 3.2 Detection of unintended edits in *APOE* Exon 4

We verified integrity and expected genotypes using the Xdrop™ enrichment workflow for 2 of the 5 cell lines (the parental *APOE*-ε3/ε4and *APOE*-KO). However, in three of the five cell lines (*APOE*-ε2/ε2, *APOE*-ε3/ε3, *APOE*-ε4/ε4) the genotype could not be confirmed. In *APOE*-ε2/ε2 and *APOE*-ε3/ε3 we could detect identical unintended insertions of ∼3.4 kb to the right of rs7412of the haplotype *APOE*-ε4 (rs429358 C; rs7412 C) (Fig. 3a). The unintended insertion caused a 5 bp deletion at positions 44,908,823 - 44,908,827 in chromosome 19 that was replaced by a 3,427 bp fragment. In the *APOE*-ε4/ε4 cell line a ~4.4 kb kb insert was identified to the left of rs429358 of the haplotype *APOE*-ε3 (rs429358 T; rs7412 C) (Fig. 3a). The unintended insertion caused a 5 bp deletion at position chr19:44,908,682-44,908,686 in chromosome 19, that was replaced by a 4,359 bp fragment. All inserts appeared to be fragments belonging to a custom pEasy Flox plasmid [26] modified with the insertion of a fragment of the human MAPT gene (Dr. Bjørn Holst (Bioneer), personal communication). The plasmid was originally introduced to provide the cells with a resistance for neomycin to enable selection of nucleofected cells and was not supposed to be present in the final edited cell lines, nor to be integrated. This was originally tested by PCR, but remained undetected [15].

**Figure 3.**
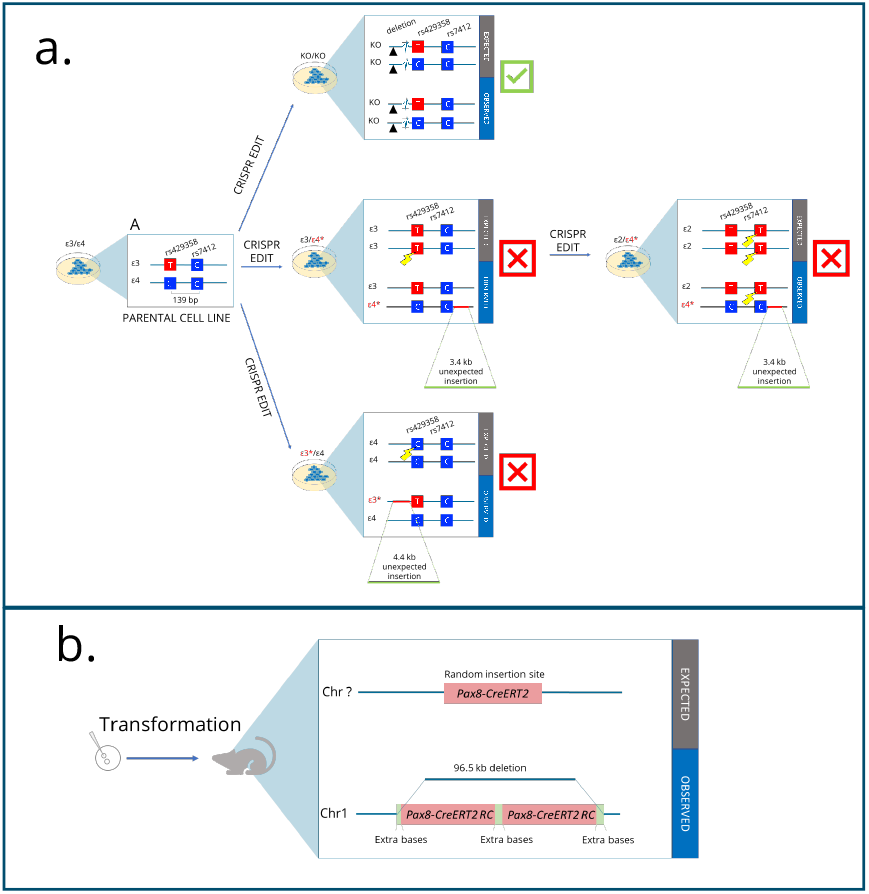
Results of the CRISPR validation by indirect sequence capture and of investigating the Pax8-CreERT2 transgene integration site in mouse. a. Overview of the resolved intended and unintended editing that occurred in the iPS cell lines. Expected and observed editing is shown for each cell line. Black triangle indicates the intended 8 bp knock-out deletion in Exon 2. Lightning indicates where editing was expected and observed. Red lines show where insertions were found with indication of insert size below. b. Overview of the identified transgene insertion site on Chr. 1 of the mouse line including the 96.5 kb deletion.

The cell line *APOE*-ε3/ε3 (initially genotyped as rs429358 T/T; rs7412 C/C upon cell line creation) was constructed from the original cell line BIONi010-C *APOE*-ε3/ε4 (rs429358 T/C; rs7412 C/C), while cell line *APOE*-ε2/ε2 (initially genotyped as rs429358 T/T; rs7412 T/T upon cell line creation) was constructed from *APOE*-ε3/ε3. It seems plausible that the observed identical insertions in *APOE*-ε3/ε3 and *APOE*-ε2/ε2 cell lines first occurred in the haplotype *APOE*-ε4 (rs429358 C; rs7412 C) when editing *APOE*-ε3/ε3 from *APOE*-ε3/ε4, and afterwards this was transmitted to the derived *APOE*-ε2/ε2 The insertion in *APOE*-ε4/ε4 appears to be a “de novo” insertion which occurred when attempting to edit the *APOE*-ε3 haplotype of the parental cell line *APOE*-ε3/ε4.

Interestingly, these unintended insertions and unedited SNPs remained undetected in the original study [15] reporting on the generation of the set of cell lines and verifying the performed editing by several complementary methods. This probably occurred because the desired editing was to create two homozygous SNPs so failure in the ability to analyse one of the two alleles would suggest that both SNPs were indeed homozygous. In all cell lines where insertions occurred, these did in fact only affect one of the two haplotypes, impairing the detection of it but leaving the other haplotype intact to be detected. In the undetected haplotype, the insert caused the amplicon encompassing the two SNPs to increase from 227 bp to > 3.4 kb or >4.4 kb, most likely hampering the amplification of the affected haplotype under the given PCR amplification conditions due to the extended length. In consequence, the edited *APOE*-ε3/ε3 cell line appeared to have the correct rs429358 T/T and rs7412 C/C genotype, the *APOE*-ε2/ε2 cell line appeared to have the correct rs429358 T/T and rs7412 T/T genotype, and the *APOE*-ε4/ε4 cell line appeared to have the correct rs429358 C/C and rs7412 C/C genotype. In reality, both the *APOE*-ε3/ε3, *APOE*-ε2/ε2, and *APOE*-ε4/ε4 cell lines kept an unedited haplotype (from the parental clone (fig. 3a). To confirm the unintended insertion, we performed sequencing of four different PCR amplicons designed to cover the region including the breakpoints on each side of the insert. Results confirmed the unintended insert in both cell lines (See supplementary Fig. 3).

### 3.3 Identification of transgene integration site in the mouse line

Next, we turned to the long-read data generated for the transgenic mouse line to see if we could identify the *Pax8-CreERT2* integration site. A total of 2544 reads were found to map to the full construct including the flanking arms with mouse *PAX8* genomic sequences located on Chr. 2. The reads mapping to the construct were mapped back to the mouse genome to identify reads that would span the breakpoint between insert and genome. Of the resulting 2544 reads, 2032 reads mapped primary to sequences shared between the construct and the mouse genome (primarily *PAX8* promotor sequences on Chr.2), but additional 45 reads mapped primary at Chr. 1 and showed the two borders of the insertion of the construct into mouse Chr. 1: 93,575,236 and at Chr1:93,671,773. Our data indicated a tandem insertion and a 96.5 kb deletion (Fig. 3b and supplementary Fig. 4). The identified insert site was confirmed by PCR across left and right breakpoints between insert and genome followed by Sanger sequencing (supplementary Fig. 5).

## 4. Discussion

In this study we have demonstrated that standard PCR-based procedures for assessing CRISPR-Cas9 genome editing events do not provide a sufficiently accurate verification. A validation of CRISPR editing should be based on examination of a larger region and the design of PCR assays should avoid designing PCR primers in the immediate vicinity of the gene editing site where occurrence of indels are frequent. The indirect sequence capture method we apply here rely on primers designed at distance 5-10 kb away from the gene editing site which provide sequence read coverage across a region around 100 kb long. This allows thorough investigation of potential unintended editing in the region surrounding the gene editing site. We have demonstrated how this technology also enable identification of transgene insertion sites with no prior knowledge on integration.

All four CRISPR-Cas9 edited cell lines analysed in our study were originally tested thoroughly by standard characterization approaches for validation of CRISPR gene editing, providing evidence of successful editing [15]. Three of these cell lines were edited at single nucleotide positions in Exon 4 to modify genotypes at two different SNPs (rs429358 and rs7412), associated with increased risk for sporadic AD. A fourth cell line was edited with an 8 bp deletion in Exon 2. Schmid et al. [15] used six complementary technologies to verify correct gene editing, including genotyping and locus integrity assessment. Genotyping relied on Sanger sequencing of a short amplicon generated across the two Exon 4 SNPs a 139 bp apart as well as another short amplicon generated across the deletion in Exon 2 in the KO cell line [15]. In the analysed mouse line, identification of transgene insertion site is made difficult by the construct flanking arms containing several kb of mouse genomic DNA and the inability to know the integrity of these at the integration site. To identify the integration site with NGA will require costly deep sequencing to obtain sufficient coverage across the breakpoints and mapping algorithms will need adjustments not to disregard unmappable breakpoint reads.

The ability to perform both long- and short-read sequencing was an advantage in our study both for the CRISPR edited cell lines and the mouse line, and should be in all instances where breakpoints are sought and larger rearrangements have occurred. It has previously been suggested that CRISPR-Cas9 editing procedures can induce complex rearrangements, deletions and chromosomal truncations around the target region [8,9,27]. These unintended events may occur as a consequence of both HDR and NHEJ repair pathways induced after CAS9 generated DSBs. The extent of these unintended events may be more substantial when CAS9 is active for extended periods of time as in plasmid based CAS9 delivery to cells but also occur with RNP- and mRNA-based delivery methods [7,27–29]. Failure to detect unintended editing can have significant consequences, especially affecting health and pharmaceutical related fields, such as pre-clinical model generation and gene therapy but also in basic research where undetected rearrangements may lead to erroneous conclusions drawn from experiments performed with edited cells.

In Kosicki et al. [8], long-range PCR amplicons followed by long read nanopore sequencing was important to show deletions several kb in size in the immediate vicinity of the site of gene editing. However, the development and optimization of long-range-PCR can be time consuming and occasionally impossible.

Other approaches have been developed for targeted enrichment of long DNA fragments e.g.:CRISPR-Cas9 systems, where selected loci are identified using guide RNA [30,31,32], and Region Specific Extraction (RSE), where a large selection of oligos is used for capturing target molecules [33]. Both these techniques can take advantage of long-read sequencing [34–36] allowing for characterization of GC-rich regions and complex genomic regions including structural variants, insertion and deletions, as well as gene duplications, repetitive regions, haplotype phasing *and de novo* genome assembly [37–39]. However, the CRISPR-Cas9 system for targeted enrichment requires relatively high amount of DNA input in the order of micrograms and results in moderate enrichment and coverage of the target region [30,31]. RSE requires design of several primers to cover larger regions and the enrichment is only around 100-fold [32]. Targeted locus amplification (TLA) is not compatible with long read sequencing but will provide locus sequence information extending over tens of kb’s.

TLA is based on physical crosslinks in adjacent sequences [40] and therefore best implemented on intact cells whereas direct analysis on isolated DNA is less straightforward. Input requirements are in the range ∼5 million cells or 5µg DNA. In addition, TLA is a relatively more elaborate and expensive assay to implement [13].

Here we have applied a new microfluidic approach for enrichment (100 to > 1000-fold) of target regions of ~ 100 kb in size, based on an indirect sequence capture approach [41]. The enriched DNA is compatible with both short and long read sequencing, which provide a broad overview of a gene edited region. In our study, this allowed us to identify and disentangle an unintended insertion that was completely missed with standard verification procedures [15]. With a slightly different primer design strategy, we were likewise able to determine the integration site for a transgene inserted with classical transformation procedures. The technology requires less than 10 ng of input DNA and only design of a single standard primer pair, which offers flexibility when targets frequently change e.g. in different gene editing experiments. Given the indirect sequence capture approach, the primer set can be designed at kilobases away from the gene editing site avoiding even kb large indels. This provides a valuable advantage in detecting large deletions, which can easily be overlooked by conventional PCR screening strategies. Especially considering that the two alleles routinely harbor deletions of distinctly different sizes [7].

### Tips and Tricks

- Recommended input material for the Xdrop™ workflow is HMW DNA (> 50 kb) of high purity (260/280 and 260/230 ratios of ~1.8 and ~2.0, respectively).
- Bead purification of the HMW DNA sample can reduce noise and improve identification and sorting of positive droplets.
- Detection Regions can be combined in multiplex reactions to extend the breadth of coverage of the Region of Interest.
- Several online tools for primer design, input calculation and enrichment assessment are available at <u>Xdrop homepage</u>.
- Simultaneous design and test of several primer sets allows to choose for the best performing based on qPCR results.
- In evaluating the performance of primers, specificity is more important than efficiency as long as good signal generation is achievable.
- The order and modality of loading reagents in Xdrop™ cartridges is very important, to ensure a successful workflow.
- Low binding plastic tubes help avoiding DNA loss, which is particular important for those steps where material is limited, such as when handling sorted positive droplets.
- It is important to avoid contamination (i.e. working in a laminar flow hood, using filtered pipette tips, etc.) when setting up the dMDA reaction. The dMDA will amplify the DNA from a single cell into > 1µg of DNA. Sorted positive droplets only contain in the fmol range of DNA so just a minor contamination will compromise enrichment.
- After sorting the positive droplets, it is crucial to keep samples on ice and proceed to the dMDA setup within 8 hours.

## Acknowledgements

This project has received funding from the European Union’s Horizon 2020 research and innovation programme under grant agreement No 848497. We would like to thank Dr. Bjørn Holst and Dr. Benjamin Schmid from Bioneer A/S for their help in data interpretation. This project has been funded in whole or in part with Federal funds from the National Cancer Institute, National Institutes of Health, under Contract No. HHSN261200800001E. The content of this publication does not necessarily reflect the views or policies of the Department of Health and Human Services, nor does mention of trade names, commercial products, or organizations imply endorsement by the U.S. Government. The authors thank Dr. Serguei Kozlov (Center for Advanced Preclinical Research) for modification of the Pax8-ERT2 transgene.

## Supplementary data

The reconstructed *APOE* reference (Chr19:44,900,000-44,920,000 including pEasy Flox fragment) has been submitted as Mendeley Data. Sequencing reads (Illumina and Oxford Nanopore) for cell lines *APOE-ε2/ε2 and APOE-ε3/ε3* mapping to the reconstructed reference have been submitted to the Genome Sequence Archive (GSA) database under the BioProject PRJCA002649.

## Appendix A

Additional equipment, consumables and reagents needed.

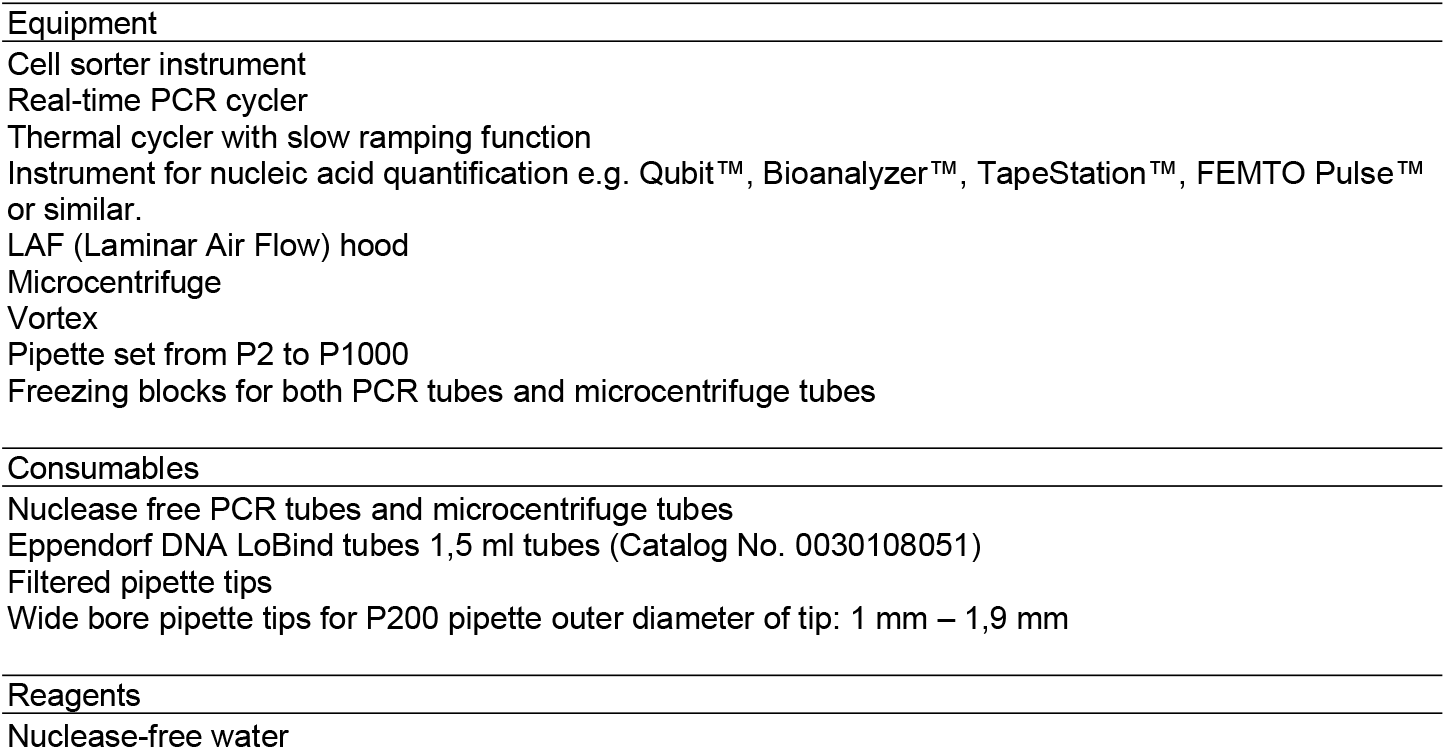

## Notes

### Competing Interest Statement

The authors have declared no competing interest.

### Summary of Updates

We have included new data on the identification of a transgene insertion site in a mouse cell line and added relevant authors. Figures have been replaced with improved versions. We have also included a more thorough comparison to alternative technologies based on recent literature references.

