## Supplementary Figures 1-5 for "Verification of CRISPR editing and finding transgenic inserts by Xdrop™ Indirect sequence capture followed by short- and long- read sequencing"

### Supplementary Figure 1

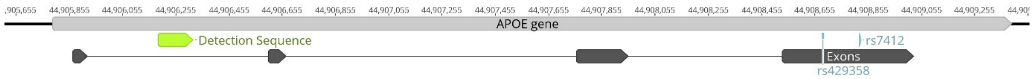


Supplementary Figure 1. Location of detection sequence in Intron 1 of the *APOE* gene used for enrichment of the gene in CRISPR edited iPS cell lines.

#### Supplementary Figure 2

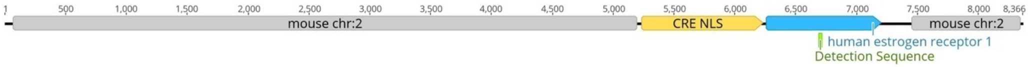


Supplementary Figure 2. The Pax8-CreERT2 transgene construct. Primer sets for detection and validation sequences were designed with the human estrogen receptor part (green).

### Supplementary Figure 3

APOE-ε3/ε3

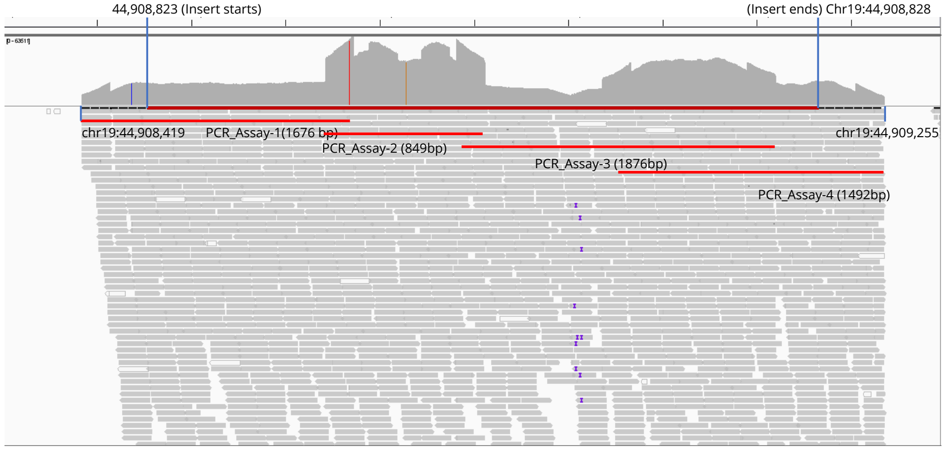


APOE-ε2/ε2

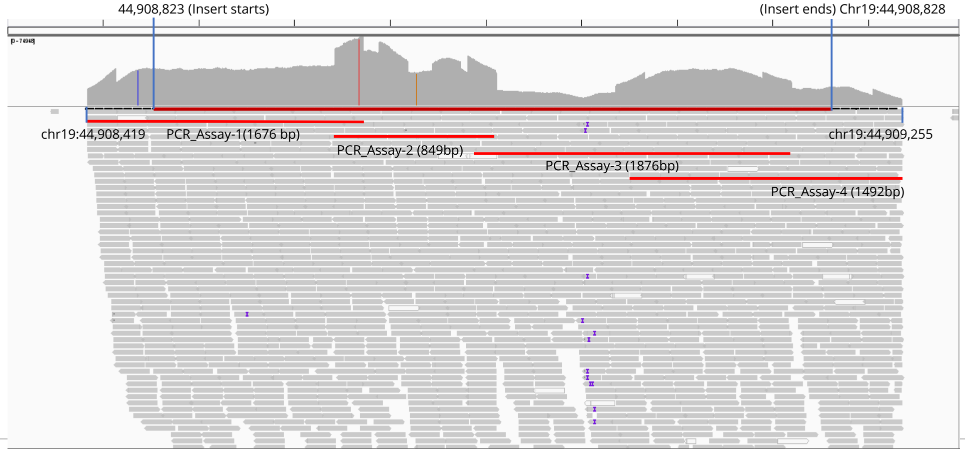


Supplementary Figure 3. Validation of APOE locus assessment made after indirect sequence capture and analysis of sequencing data. Short-read sequencing coverage plots (IGV) from the PCR validation of the 3.4 kb pEasyFlox insert in APOE-ε3/ε3 and APOE-ε2/ε2. PCR products 1-4 are marked with sizes on the plot as well as the insert borders in the Chromosome 19 and the PCR validation fragments. The insert was validated in both cell lines.

#### Supplementary Figure 4

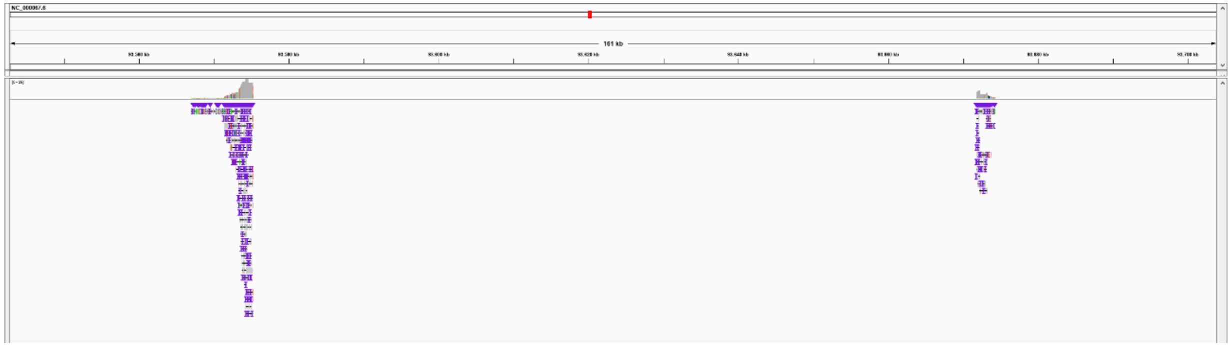

Supplementary Figure 4. Long reads identified to map to the Pax8-CreERT2 transgenic construct and Chr. 1 and indicating the left and right borders between insert and Chr. 1. In addition, a 96.5 kb deletion is indicated.

### Supplementary Figure 5

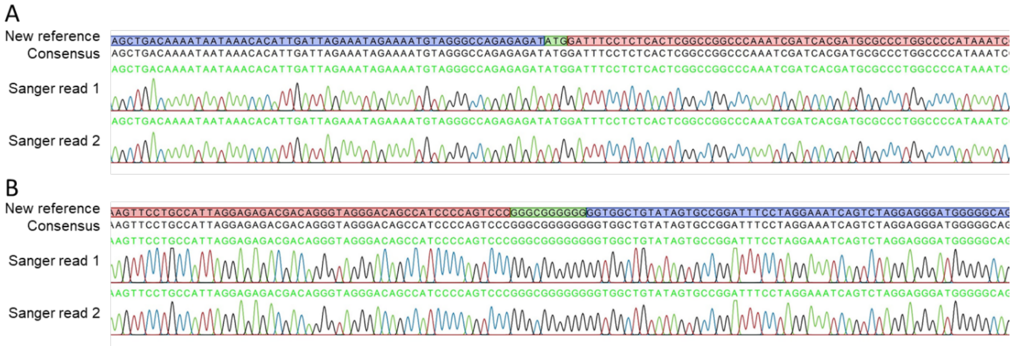


Supplementary Figure 5. **Validation of Pax8-CreERT2 insertion site in the transgenic mouse line.** Sanger sequencing of two different PCR products spanning the left and right border, respectively. The construct (red) was inserted reverse complement into chromosome 1 (blue) and additionally on left and right border 3 and 10 extra bases (green) were inserted, respectively.
